# Latent Gaussian Process Modeling for Dynamic PET Data: A Hierarchical Extension of the Simplified Reference Tissue Model

**DOI:** 10.64898/2026.04.11.717906

**Authors:** Johan Vegelius

**Affiliations:** Department of Medical Sciences, Experimental Cognitive and Affective Neuroscience Lab, Uppsala University, Uppsala, Sweden; Department of Psychology, Uppsala University, Uppsala, Sweden

**Keywords:** hierarchical model, positron emission tomography, simplified reference tissue model, Gaussian process

## Abstract

Dynamic positron emission tomography (PET) provides a powerful tool for studying in vivo neurochemical processes, including transient neurotransmitter release. However, widely used models such as the simplified reference tissue model (SRTM) assume time-invariant kinetic parameters, limiting their ability to capture dynamic changes in binding. Existing extensions introduce time-varying effects through parametric response functions or basis expansions, but are often constrained by restrictive functional assumptions, computational complexity, or limited uncertainty quantification.

We propose a latent Gaussian process extension of SRTM (LGPE-SRTM), in which the apparent efflux parameter is modeled as a smooth, time-varying function within a hierarchical framework. By applying a first-order implicit discretization of the governing differential equation, the model admits a representation that is linear in all kinetic parameters in the mean structure, while nonlinearity is confined to a parameter-dependent covariance. This yields a conditionally linear mixed-effects model with structured covariance, enabling efficient likelihood-based inference.

The proposed approach integrates mechanistic modeling, nonparametric flexibility, and hierarchical inference in a unified and computationally scalable framework. By representing functional effects on a shared temporal domain, all core computations reduce to operations on low-dimensional matrices whose size is independent of the number of subjects. This enables robust population-level inference on timevarying neurotransmitter dynamics without imposing restrictive parametric forms.

The method is demonstrated on both simulated and empirical PET data, where it accurately recovers transient effects, provides well-calibrated uncertainty, and distinguishes constant from time-varying dynamics.

## 1. Introduction

Dynamic positron emission tomography (PET) provides a powerful tool for studying in vivo neurochemical processes, including transient neurotransmitter release. A widely used approach for quantifying receptor binding is the simplified reference tissue model (SRTM), which avoids arterial sampling by using a reference region (Lammertsma & Hume, 1996). In differential form, SRTM can be written as

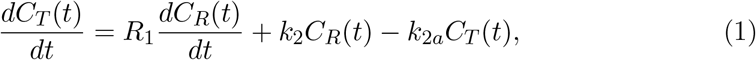

where *C*_*T*_ (*t*) and *C*_*R*_(*t*) denote tracer concentrations in the target and reference regions, respectively, and (*R*_1_, *k*_2_, *k*_2*a*_) are kinetic parameters governing tracer delivery and efflux. A key limitation of SRTM is the assumption of time-invariant parameters, which precludes modeling transient changes in binding during the scan.

Several extensions have been proposed to address this limitation. The linear simplified reference region model (LSRRM) (Alpert et al., 2003) introduces time-varying binding through a parametric perturbation of *k*_2*a*_(*t*), retaining linear estimation at the cost of restrictive assumptions on the temporal response. More flexible parametric approaches, such as ntPET (Morris et al., 2005), employ biologically motivated response functions but require nonlinear estimation, which can be computationally demanding and sensitive to initialization. Linearized variants, including lp-ntPET (Normandin et al., 2012), approximate the transient response using a finite basis expansion, improving computational tractability but constraining the dynamics to a predefined functional space.

Nonparametric approaches (C. Constantinescu et al., 2008; C. C. Constantinescu et al., 2007) allow the transient signal to vary freely over time, typically through regularized inverse problems. While increasing flexibility, these methods are inherently ill-posed, with the estimated trajectory largely determined by regularization rather than the data, and do not provide a fully probabilistic characterization of uncertainty.

Bayesian extensions, (e.g., Irace et al., 2020) offer principled uncertainty quantification but are generally restricted to low-dimensional parametric representations and are computationally challenging in hierarchical settings. More recent likelihood-free approaches such as PET-ABC (Fan et al., 2021) avoid explicit likelihood evaluation but may suffer from low statistical efficiency and do not directly yield explicit estimates of time-varying functions, particularly in flexible nonparametric form.

A further limitation of existing approaches is the treatment of measurement noise. Most models assume independent errors across time frames, despite the fact that discretization of the underlying compartmental system induces temporal dependence through noise propagation. Ignoring this structure leads to misspecified likelihoods and potentially miscalibrated uncertainty quantification.

In this work, we propose a latent Gaussian process extension of SRTM (LGPE-SRTM) that addresses these limitations within a unified framework. The key idea is to model the apparent efflux parameter *k*_2*a*_(*t*) as a latent function governed by a Gaussian process within a hierarchical mixed-effects formulation. By discretizing the governing differential equation using a first-order implicit scheme, the model admits a representation that is linear in all kinetic parameters in the mean structure, while nonlinearity is confined to a parameter-dependent covariance arising from propagated measurement noise. A key obstacle to hierarchical functional PET modeling is that computational cost scales with the total number of observations. We show that, by restructuring the model on a shared temporal domain, all core computations reduce to *M × M* matrices, where *M* is small and independent of the number of subjects.

This formulation yields several advantages. First, it combines mechanistic modeling with nonparametric flexibility, allowing data-driven estimation of time-varying binding without imposing restrictive parametric forms. Second, the hierarchical structure enables partial pooling across subjects, improving statistical efficiency and supporting population-level inference. Third, explicit modeling of the induced temporal covariance ensures coherent uncertainty quantification. Finally, computational tractability is achieved by representing all functional effects on a shared domain via the matrices **E**_*j*_, which map subject-specific observations to a common *M*-dimensional space. This reparameterization reduces all core computations to operations on *M × M* matrices, where *M* is the number of unique time points at which the time-varying effect is defined, typically on the order of tens. Crucially, *M* does not grow with the number of subjects *N*. As a result, the dimension of the key precision matrix **Δ**, with *X*_*j*_ and *V*_*j*_ being the subject-specific design matrix and covariance matrix, respectively, and 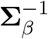 is the prior precision matrix of the fixed effects, then

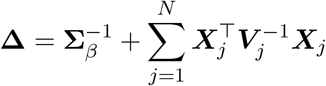

remains fixed as *N* increases, and only its components are updated additively across subjects. This implies that the computationally dominant step (matrix inversion) scales with *M* rather than with the total number of observations. In contrast, standard nonparametric or hierarchical formulations operate in the full observation space, where matrix dimensions scale with Σ_*j*_ *m*_*j*_, (*m*_*j*_ being the number of observations per subject) leading to cubic computational complexity in the number of observations. By decoupling the functional representation from the observation space, the proposed approach achieves substantial computational savings and enables scalable inference in large cohorts without sacrificing model flexibility. Unlike lp-ntPET, which restricts dynamics to a finite basis, and nonparametric inverse methods, which are ill-posed, the proposed model achieves flexible functional estimation with identifiable and probabilistically coherent inference.

Together, these features bridge a gap between mechanistic compartment modeling, nonparametric function estimation, and hierarchical statistical inference, providing a flexible and scalable framework for analyzing dynamic PET data with timevarying neurotransmitter effects.

Related hierarchical functional modeling approaches have also been proposed by the author in other application domains (Hoppe et al., 2025, e.g.,), where time-varying effects are modeled using Gaussian process-based mixed models and estimated via marginal likelihood with empirical Bayes updates. In that work, the framework is applied to longitudinal clinical data without mechanistic constraints. The present work extends these ideas to dynamic PET by incorporating a compartmental model and an induced covariance structure arising from the discretized system.

## 2. Method

### First-order discretization

In the proposed extension, the apparent efflux parameter *k*_2*a*_ is allowed to vary smoothly over time. For subject *j* = 1, …, *N*, the model is

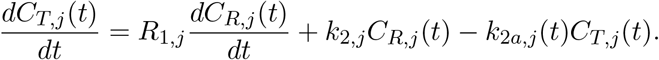

Let 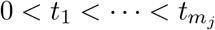 denote observation times and define Δ*t*_*m*_ = *t*_*m*_ − *t*_*m*−1_. Using a first-order implicit (backward Euler) discretization yields

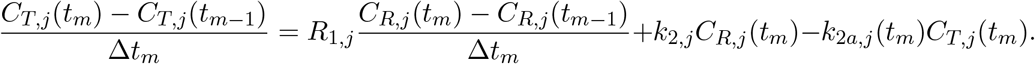

Let *m*_*j*_ ∈ ℕ^+^ be the number of times at which *C*_*T,j*_ and *C*_*R,j*_ are observed. Define vectors 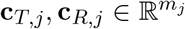 and the differencing matrix 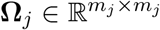. Then

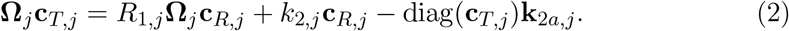

### Functional random effects via Gaussian processes

Let 𝒮 = *{s*_1_, …, *s*_*M*_ *}* be a common domain defined via a transformation *s* = *f* (*t*), e.g. *s* = max(0, *t* − *T*_*c*_), allowing *k*_2*a*_ to vary only after an intervention time *T*_*c*_.

We model

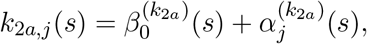

where

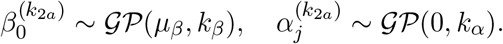

A typical choice is the squared exponential kernel

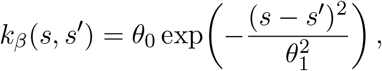

where *θ*_0_ controls variance and *θ*_1_ the length scale. In the empirical and simulated data in this work, squared exponential kernel was used. Discretizing on 𝒮, we obtain

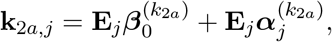

where **E**_*j*_ maps the common grid to subject-specific times. Hence, ***E***_*j*_ is a *m*_*j*_ *× M* - matrix where *m*_*j*_ is the number of measurements of subject *j* and *M* is the number of unique times of observations across the sample where *k*_2*a*_ is allowed to vary. This is particularly convenient when we assume that *k*_2*a*_ is fixed before an intervention and varies afterward. In addition missing values are handled automatically via the ***E***_*j*_ matrix. Scalar parameters are modeled hierarchically:

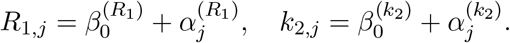

Kinetic parameters in compartment models, including *R*_1_, *k*_2_, and *k*_2*a*_, are known to exhibit substantial correlation (Matheson & Ogden, 2022). In the present work, the primary focus is on the functional form of *k*_2*a*_(*t*), and a simplified covariance structure is therefore adopted. While a fully joint covariance structure could in principle be specified, it is expected to be weakly identifiable in typical dynamic PET settings with limited temporal resolution. Estimating such a structure would introduce additional uncertainty without clear gains for inference on the time-varying behavior of *k*_2*a*_(*t*). The independence assumption thus acts as a pragmatic regularization that enables stable estimation of the functional component of primary interest. Assuming independence across components,

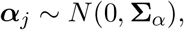

contain variances and covariances across both scalar (*R*_1,*j*_ and *k*_2_) and functional (*k*_2*a*_) parameters with block-diagonal covariance.

The Gaussian process length-scale parameter was fixed at *θ*_1_ = 10 minutes, corresponding to the expected temporal scale of neurotransmitter dynamics and the effective resolution of the PET signal. In this setting, the length-scale is weakly identified due to the limited number of observation points, and attempts to estimate it jointly with other model components led to unstable behavior. Fixing *θ*_1_ therefore provides a principled regularization that stabilizes inference without materially restricting the class of admissible trajectories.

For identifiability and stability, the signal variance parameter of the Gaussian process (*θ*_0_) was constrained to be equal across the intercept and group-level functions. This reflects the fact that both components represent variation on the same biological scale (the apparent efflux parameter *k*_2*a*_(*t*)), while reducing the number of weakly identifiable hyperparameters in the model.

### Measurement model and induced autocorrelation

Observed TACs satisfy

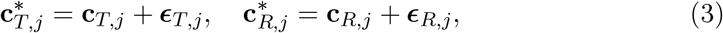

with

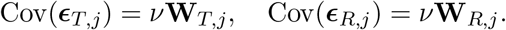

Now define

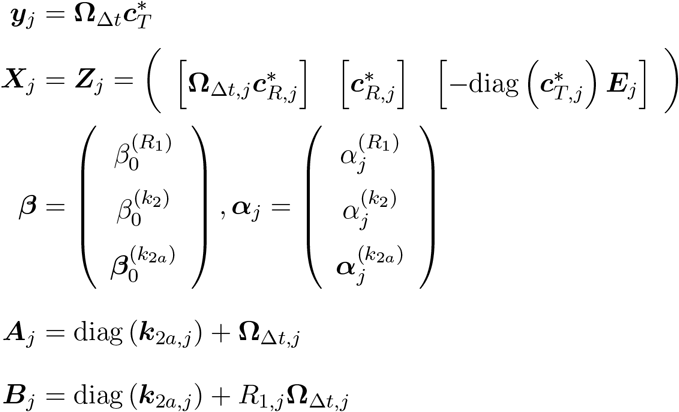

Using (3) and substituting into (2) yields

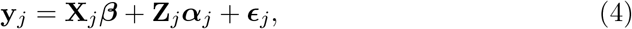

with

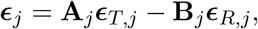

and covariance

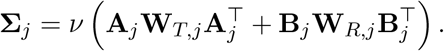

This yields a linear mixed model with parameter-dependent covariance, where all nonlinearity is confined to Σ_*j*_.

### Marginal model

Integrating out ***α***_*j*_ yields

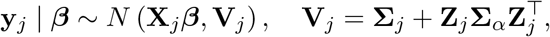

where **Σ**_*α*_ is the prior distribution of ***α***_*j*_ obtained from the kernel. Assuming a Gaussian prior ***β*** ∼ *N* (***µ***_*β*_, **Σ**_*β*_), the conditional posterior distribution is

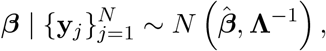

where

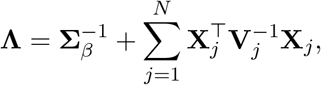

and

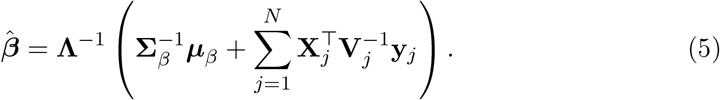

Notice that **Λ** is an *M × M* -matrix, independent of *N*.

### Estimation

Estimation is performed using an iterative scheme alternating between: (i) updating ***β*** via Equation (5) (including uncertainty via **Λ**), (ii) updating ***α***_*j*_ via best linear unbiased prediction (empirical Bayes posterior means), (iii) updating subject-specific kinetic parameters, which enter the model through the covariance matrices **Σ**_*j*_ and are treated as plug-in estimates, (iv) updating variance components based on the current random effects estimates, and (v) optimizing kernel hyperparameters via marginal likelihood, where the fixed effects ***β*** are integrated out.

The tri-diagonal structure of **Σ**_*j*_ enables computationally efficient implementation. The procedure combines marginal likelihood estimation for fixed effects and covariance parameters with empirical Bayes updates of subject-specific random effects.

## 3. Illustration

We demonstrate the proposed LGPE-SRTM on both empirical and simulated dynamic PET data. The results are shown in Figure 1.

**Figure 1:**
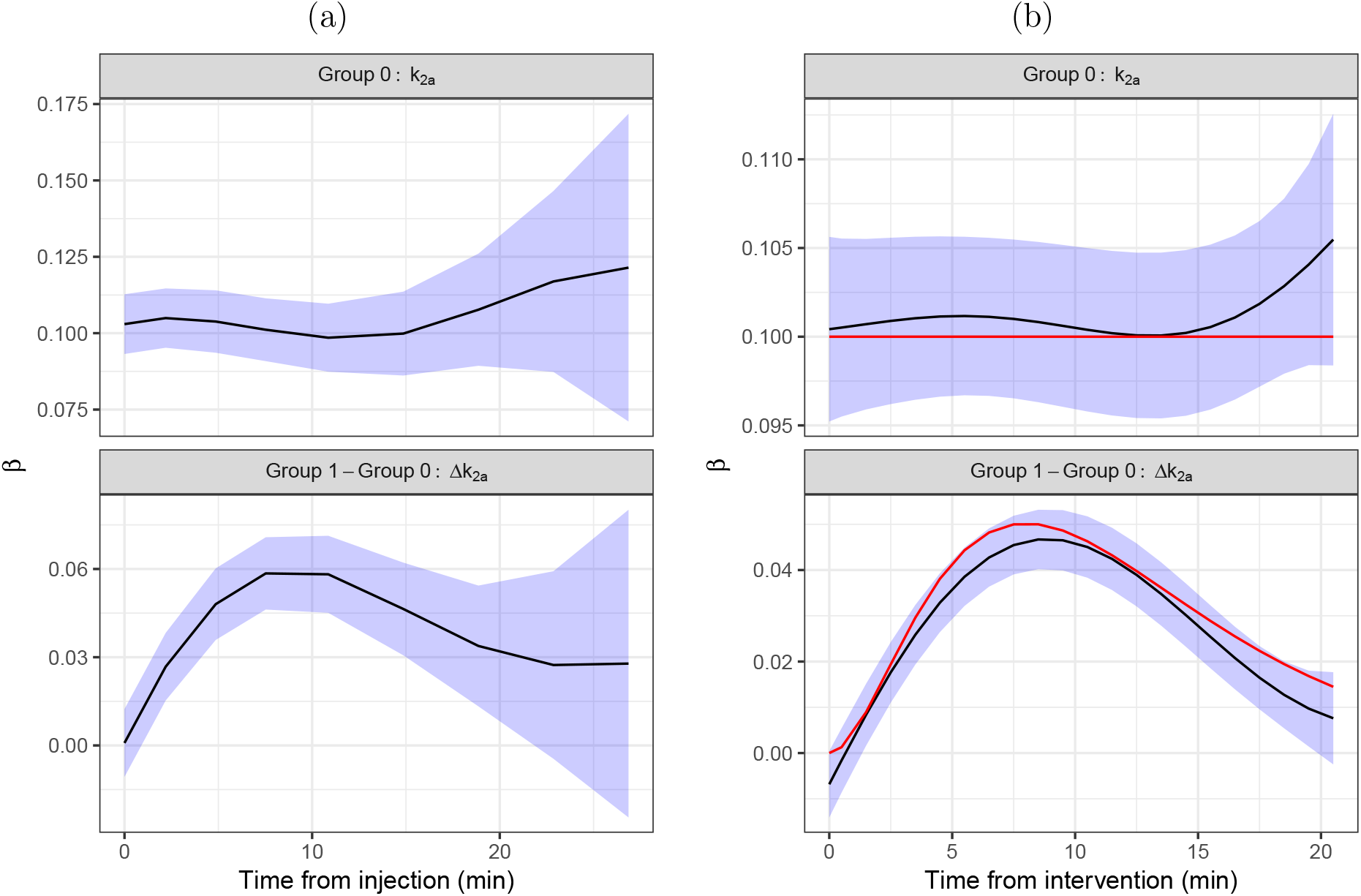
Estimated time-varying apparent efflux parameter *k*_2*a*_(*t*) and group differences Δ*k*_2*a*_(*t*) from empirical and simulated data. The left panel shows results from the empirical data set (Johansson et al., 2024) (*N* = 20), with the saline group (top) and the difference between amphetamine and saline groups (bottom). The right panel shows results from a simulated data set (*N* = 50), including the true underlying trajectories (red curves). Black curves denote estimated population-level functions and shaded regions represent 95% pointwise uncertainty bands. In both empirical and simulated data, no significant time-varying effect is detected in the reference group under the null hypothesis of a constant *k*_2*a*_(*t*), whereas significant time-varying effects are detected for group differences. In the simulated setting, the estimated trajectories closely recover the true underlying functions. These results demonstrate that the proposed LGPE-SRTM provides well-calibrated uncertainty and reliably distinguishes constant from time-varying functional effects.

The empirical data (Johansson et al., 2024) consist of *N* = 20 rodents, of which 14 received an amphetamine injection and 6 received a saline (placebo) injection after approximately 25 minutes (left panel). The estimated intercept function, corresponding to the *k*_2*a*_(*t*) trajectory in the saline group, showed no evidence of time variation following injection (*χ*^2^(edf = 2.87) = 2.74, *p* = 0.41), where edf represents the effective degrees of freedom (Wood, 2013), consistent with the null hypothesis of a constant trajectory. In contrast, the estimated difference function between amphetamine and saline groups revealed a clear time-varying effect (*χ*^2^(edf = 2.75) = 123.36, *p <* 0.001), indicating a transient increase in *k*_2*a*_(*t*) following amphetamine administration.

A similar pattern was observed in the simulated data (right panel, *N* = 50), where the true underlying trajectories were known. For the reference group with constant *k*_2*a*_(*t*), no deviation from constancy was detected (*χ*^2^(edf = 3.71) = 3.07, *p* = 0.41). In contrast, the group difference exhibited a strong time-varying effect (*χ*^2^(edf = 3.49) = 1036.61, *p <* 0.001), closely matching the true underlying trajectory.

Across both empirical and simulated settings, the method correctly identifies null (constant) trajectories as non-significant while detecting significant deviations for time-varying effects. In the simulated data, the estimated functions closely recover the true temporal dynamics, both in shape and magnitude. These findings support the validity of the proposed framework and demonstrate its ability to provide reliable inference on time-varying neurotransmitter effects without relying on parametric assumptions.

## 4. Limitations

The proposed approach relies on a first-order discretization of the underlying kinetic system, which introduces approximation error that may become non-negligible when observation intervals are large or when the true dynamics exhibit rapid temporal variation. The use of Gaussian process priors provides flexibility but introduces sensitivity to kernel choice and hyperparameter estimation, which can affect the identifiability and smoothness of the estimated functions, particularly in low signal-to-noise regimes. Although the hierarchical structure enables partial pooling and improved statistical efficiency, disentangling time-varying effects from noise remains challenging in settings with limited data or weak signal. Furthermore, while the computational burden is substantially reduced through the common-domain reparameterization, extensions to multi-region or high-dimensional settings may still require careful algorithmic optimization

In the current implementation, the Gaussian process length-scale was fixed and the signal variance parameter was constrained to be equal across functional components to ensure stable estimation. These restrictions act as regularization in a setting where hyperparameters are weakly identifiable given the limited temporal resolution of the data. Future work will consider fully data-driven approaches, including fully Bayesian formulations in which hyperparameters such as *θ*_0_ and *θ*_1_ are assigned prior distributions.

A systematic comparison with existing approaches, including lp-ntPET and models that ignore induced temporal dependence, is an important direction for future work. This will clarify the extent to which explicitly modeling time-varying kinetics and error structure improves inference and uncertainty calibration in practice.

